# Interactive computational and experimental approaches improve the sensitivity of periplasmic binding protein-based nicotine biosensors for measurements in biofluids

**DOI:** 10.1101/2023.01.16.524298

**Authors:** Nandan Haloi, Shan Huang, Aaron L. Nichols, Eve J. Fine, Nicholas J. Friesenhahn, Christopher B. Marotta, Dennis A. Dougherty, Erik Lindahl, Rebecca J. Howard, Stephen L. Mayo, Henry A. Lester

## Abstract

We developed fluorescent protein sensors for nicotine with improved sensitivity. For iNicSnFR12 at pH 7.4, the proportionality constant for ΔF/F_0_ vs [nicotine] (δ-slope, 2.7 μM^-1^) is 6.1-fold higher than the previously reported iNicSnFR3a. The activated state of iNicSnFR12 has a fluorescence quantum yield of at least 0.6. We measured similar dose-response relations for the nicotine-induced absorbance increase and fluorescence increase, suggesting that the absorbance increase leads to the fluorescence increase via the previously described nicotine-induced conformational change, the “candle snuffer” mechanism. Molecular dynamics (MD) simulations identified a binding pose for nicotine, previously indeterminate from experimental data. MD simulations also showed that Helix 4 of the periplasmic binding protein (PBP) domain appears tilted in iNicSnFR12 relative to iNicSnFR3a, likely altering allosteric network(s) that link the ligand binding site to the fluorophore. In thermal melt experiments, nicotine stabilized the PBP of the tested iNicSnFR variants. iNicSnFR12 resolved nicotine in diluted mouse and human serum at 100 nM, the peak [nicotine] that occurs during smoking or vaping, and possibly at the decreasing levels during intervals between sessions. NicSnFR12 was also partially activated by unidentified endogenous ligand(s) in biofluids. Improved iNicSnFR12 variants could become the molecular sensors in continuous nicotine monitors for animal and human biofluids.

## Introduction

The iNicSnFR series of intensity-based fluorescent nicotine-sensing fluorescent reporters was initially developed to test the hypothesis that nicotine enters the cytoplasm and organelles at the concentrations relevant to the brain of smokers and vapers [1]. This entry allows nicotine to act as a pharmacological chaperone for nascent nicotinic acetylcholine receptors in the endoplasmic reticulum. This process leads to the upregulation thought to play a key role in nicotine dependence [2]. Previous pharmacokinetic measurements on the blood of smokers and vapers indicate that the peak of [nicotine] in the blood is 100-200 nM; and in the interval between puffing sessions, [nicotine] declines to concentrations as low as 10 nM [3]. The initial sensors, iNicSnFR3a and iNicSnFR3b, lack the sensitivity for measurements in this range; but the linearity of the sensors allowed us to extrapolate to lower concentrations in experiments on various cultured cell types. We therefore considered the hypothesis verified [1].

Previous pharmacokinetic measurements on the blood of smokers and vapers utilize intravenous catheters followed by HPLC- and mass spectrometry-based assays. These measurements are invasive and tedious. Our goals now include incorporating fluorescent reporters into wearable, minimally invasive and non-invasive devices that provide continuous measurement of nicotine pharmacokinetics in human interstitial fluid or sweat during smoking or vaping sessions. This application requires iNicSnFR constructs that detect nicotine at 10 nM, with a time resolution of ∼ 5 min.

Therefore, we sought to improve the sensitivity of the iNicSnFR series. Each member of the iNicSnFR series is a merger of two proteins: a suitably mutated, interrupted, OpuBC periplasmic binding protein, and a circularly permuted GFP (cpGFP). The present fluorescent sensors are related both to the acetylcholine (ACh) sensor iAChSnFR [4] and to several other sensors which we term iDrugSnFRs because the OpuBC binding site has been mutated to selectively bind individual central nervous system-acting drugs [1, 5–7]. In all cases but ACh, the drug is a weak base (6.5 < pKa < 10), and the charged form of the drug binds to the PBP. Because atomic-scale structural data are available for the entire iNicSnFR3a protein, in both apo and varenicline-bound states [6], it is now possible to test structure-based hypotheses to improve the sensitivity.

Here we combined computational protein design with mutagenesis, fluorescence, and absorbance assays. The resulting sensor, iNicSnFR12, is ∼ 6.1-fold more sensitive than previous constructs, allowing measurements in biofluids at concentrations below the 100 nM level. Although diffuse electron density in X-ray data [6](PDB 7S7T) previously vitiated an atomic-level description of nicotine-protein interactions, here we further used molecular simulations to model a ligand pose consistent with experiments, and to propose an allosteric basis for altered sensitivity in iNicSnFR12. Therefore, we report good progress toward the goal of a protein biosensor that can serve in minimally invasive devices to continuously monitor nicotine in the blood of smokers or vapers.

## Methods

### Mutagenesis and library screening

Site-saturated mutagenesis (SSM) libraries were generated employing the so-called “22c trick” [8] for PCR primers. For site-directed mutagenesis (SDM), individual primers were used for PCR. The PCR products were gel purified, digested with DpnI, ligated using the Gibson Assembly Master Mix (New England BioLabs), and used to directly transform *E. coli* BL21-Gold (DE3) chemically competent cells (Agilent Technologies, Santa Clara, CA). The expression and screening of iNicSnFR variants were performed in 96-well plate format, and 93 variants per SSM library and 186 variants per combinatorial mutagenesis (CM, see below) library were screened. Individual colonies from iNicSnFR libraries were cultivated in 1 ml of ZYM5052 autoinduction media [9] supplemented with 100 μg/ml ampicillin for 30 h at 30 °C, 250 rpm. Then the cells were harvested (3000 × g, 10 min, 4 °C), washed with PBS, pH 7.40, and resuspended in 3×PBS, pH 7.40. Resuspended cells were lysed by freezing and thawing using liquid N_2_ and a room-temperature water bath. Intact cells and cell debris were removed by centrifugation at 3500 × g for 30 min at 4°C. The supernatants of lysates were tested with excitation at 485 nm and emission at 535 nm in the presence or absence of a nicotine concentration near the EC_50_ of the parent construct for the library being screened. A Spark M10 96-well fluorescence plate reader (Tecan, Männedorf, Switzerland) was used to measure baseline fluorescence (F_0_) and nicotine-induced fluorescence (ΔF). Promising clones were amplified and sequenced, and the beneficial mutations were confirmed by the measurement of dose-response relations against nicotine (described below). The optimally responding construct in each round of mutagenesis was used as a template for the next round of SSM, SDM, or combinatorial mutagenesis (CM).

### Purification of iNicSnFRs

Proteins were expressed in *E. coli* BL21-Gold (DE3) cells using 50 ml ZYM5052 autoinduction media [9]. Cells were collected by centrifugation and stored at –80°C until use. For purification, frozen cell pellets were resuspended in PBS, pH 7.40, and lysed by sonication. Intact cells and cell debris were removed by centrifugation at 15,000 × g for 30 min at 4°C. The supernatant was collected and loaded onto a prewashed Ni-NTA column with wash buffer at 4°C. Ni-NTA wash buffer contained 20 mM imidazole in PBS, pH 7.4. Elution was achieved using an imidazole gradient (20–200 mM). Proteins were concentrated by centrifugation through Amicon Ultra 15 filter units (Millipore, Burlington, MA) with a 30-kD cutoff and then dialyzed against 3×PBS, pH 7.40. The dialyzed protein was then subjected to dose-response studies to characterize its responses to various drugs.

### Fluorescence dose-response relation measurement

Purified biosensors were mixed with drug solutions to yield the mixture of 100 nM biosensor and the drug with desired concentrations. Samples were tested in at least three wells (see individual figures below). Solutions were in 3×PBS, pH 7.0 or 7.4. A Tecan Spark 10M plate reader was used to read the plate with 485 nm excitation and 535 nm emission wavelengths to measure GFP fluorescence (F_0_ and ΔF). The resulting data were fitted with Origin 9.2 software (OriginLabs) to the Hill equation [1, 6].

### Absorbance measurements

Absorption spectra were measured in a Tecan Spark 20M instrument. Chromophore concentration was measured by denaturation in 1 M NaOH using the absorption coefficient of 44,000 cm^-1^ M^-1^ at 447 nm [10]. To a solution of iNicSnFR12 (7.3 μM in 3×PBS, pH 7.0), we added increments of the same solution with added 1 mM nicotine tartrate (Thermo Fisher, Waltham, MA). The relationship between absorption at 496 nm and [nicotine] was fitted to the molar absorptivity of the apo and bound sensors (ε_apo_ and ε_bound_, respectively) and the K_m_ for the nicotine-iNicSnFR12 complex (*K*) assuming a single binding site, corrected for depletion of the free [nicotine] by binding to iNicSnFR12:

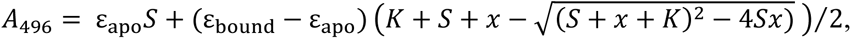

where *S* is the concentration of iNicSnFR12 and x is the total nicotine concentration.

### Quantum yield measurements

Quantum yields (Φ) were measured by comparison with the value for EGFP. As above, chromophore concentration was measured by denaturation in 0.1 M NaOH using the absorption coefficient of 44,000 cm^-1^ M^-1^ at 447 nm [10]. The uncorrected relative fluorescence quantum yield for the sample is given by [11]

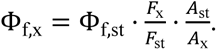

Φ_f,st_ is the fluorescence quantum yield of the reference standard, which was purified EGFP (Φ_f,st_ = 0.60 at λ_ex_ = 489 nm) [12]. *F* is the integrated fluorescence emission for 489 nm excitation. *A* is the absorbance at 489 nm. The iNicSnFR variants were measured in 3×PBS, pH 7.4 with [nicotine] = 10×EC50 for each variant. EGFP was measured in 3×PBS, pH 7.4. Background-subtracted absorption spectra for samples at protein concentrations between 3-4 μM were measured in a 10-mm pathlength cuvette in the Tecan Spark 20M instrument. Samples were then diluted 20-fold in 3×PBS, pH 7.4 with the same [nicotine] in a clear-bottom 96-well plate, and fluorescence measurements were taken by exciting at 489 nm (20 nm nominal bandwidth) with the monochromator and detecting emissions (1 nm step size; 20 nm bandwidth). Fluorescence assay volumes were 337 μL. Background-subtracted emission was integrated between 528 (to avoid overlap with the excitation) and 650 nm.

The Φ values for nicotine-saturated iNicSnFR fluorescence were corrected by three factors. Factor 1 accounts for the extrapolation of iNicSnFR values taken at 10 x EC_50_ to predicted values at saturated [nicotine]; this factor equals 1.1. Factor 2 accounts for relative integrated absorbed photons by the iNicSnFR3a, iNicSnFR11, or iNicSnfFR12 vs GFP at the peak excitation wavelength (489 nm) and effective excitation bandwidth of 26 nm, equal to 0.957. Factor 3 accounts for the shift in peak emission of iNicSnFR3a, iNicSnFR11, or iNicSnfFR12 vs that of EGFP, resulting in a different fraction of total fluorescence when we integrated from 528 to 700 nm. The emission peak for iNicSnR3a is shifted by 5 nm vs that for EGFP [1], and we noted that the emission decreases at a rate of e-fold per 31 nm. Factor 3 is therefore *e*^-5/31^ = 0.851. We corrected the observed quantum yield by the product of these factors, 0.895.

### Computational search for docking poses of nicotine

A workflow resembling a previous study [13] was developed to systematically search for the optimal binding pose of nicotine in the diffuse electron density near the binding site in the X-ray map of iNicSnFR3a (PDB: 7S7U) [6]. To reduce the search space, we used knowledge of the well-resolved varenicline-iNicSnFR3a interaction. Both ligands have a protonated amine group. Varenicline is known to form multiple cation-π interactions within the binding site of iNicSnFR3a [1, 6]; and nicotine is known to form at least one these cation-π interactions, with Tyr-357 [1]. Assuming that other cation-π interactions known for varenicline also occur for nicotine, we first aligned the protonated amine of nicotine to the secondary amine of varenicline. To optimize possible resolution and placement, we placed the pyrrolidine amine using the coordinates of the highest-resolution liganded structure in the iNicSnFR series, the complex between varenicline and the PBP from iCytSnFR (PDB: 7S7X). We defined a vector, from the pyrrolidine amine in nicotine to the C3 carbon atom of the pyridine ring. We mapped this vector onto a Fibonacci spherical lattice (FSL) [14] of 25 points, generating multiple orientations. During rotation, the amine group defined the center of the FSL, while the carbon atom was arranged on the FSL. Each of the 25 generated orientations at each grid point was then self-rotated around the vector, with an interval of Ψ = 45°, creating a total of 200 poses. To account for the internal degrees of freedom, multiple ligand conformations were generated using the Open Babel toolkit [15]. The procedure was repeated for each generated conformer, resulting in a total of 1000 poses.

During our search process, bad contacts between protein/drugs, such as clashes or piercings, are inevitable. Poses associated with clashes and ring piercings were detected [13] and removed, resulting in a total of ∼ 600 distinct poses.

Each pose was then subjected to 1,000 steps of energy minimization using the generalized Born implicit solvent (GBIS) module implemented in NAMD2 [16, 17]. GBIS calculates molecular electrostatics in solvent by representing water as a dielectric continuum as described by the Poisson-Boltzmann equation. Therefore, GBIS accounts for the effect of solvation on nicotine-protein interactions without including explicit solvent, which greatly accelerates modeling efficiency. During minimization, the protein side chains and the drug were allowed to move, while the protein backbone was kept fixed in order to allow the drug to relax in its environment without causing a significant conformational change in the protein. We chose to maintain the global conformation of the protein, because the X-ray structure of iNicSnFR3a (PDB 7S7U) is already in the nicotine-bound form.

The energy-minimized poses were then used to start molecular dynamics (MD) simulations of the liganded iNicSnFR3a (PDB 7S7T) complex with explicit water and ion neutralized system. To start simulations from structurally different states, we first clustered the poses based on root-mean square deviation (RMSD) of the drug with a cutoff of 1.5 Å and chose 10 clusters. From each cluster, a representative pose was selected (close to the centroid) and solvated with explicit water and neutralized with 150 mM KCl, resulting in a total of ∼ 120,000 atoms with the dimension of 10.9 X 10.9 X 10.9 nm. Each system was energy-minimized using the steepest-descent algorithm for 5,000 steps and then relaxed with the protein restrained, independently. The position restraints were gradually released during the first 4 ns to allow adjustment of the nicotine within the binding pocket **(Table S1)**. Then, unrestrained simulations were performed for all 10 simulations, each for 200 ns, gathering a total of 2 µs of simulations.

The effect of the nicotine binding on the protein stability was monitored in each simulation starting from the 10 poses. In seven simulations, the PBP undergoes a conformational change from the holo- to apo-like state (i.e., opening of the binding cleft), suggesting that the particular nicotine poses in these simulations are not favorable. The three simulations that showed most stability of the PBP (Figure S1) were analyzed further to model the final nicotine-bound structure. We re-clustered these simulation data and found only three major clusters with populations greater than 10%. To determine the pose that can fit best to the X-ray map, we first converted the X-ray structure factor file to a real-space map file using the *phenix.mtz2map* command in *phenix* [18] and then calculated the cross-correlation of a representative pose from each cluster (structure closed to the centroid) with that of the generated map. The pose from the highest populated cluster and with the greatest cross-correlation is chosen as the best representation of the nicotine-bound structure.

### Biofluids

Human serum was obtained from Sigma-Aldrich. Rat serum was obtained from Sigma-Aldrich. Heat treatment was for 65° C, for 10 min, followed by passage through a 0.2 μm Millipore filter.

### MD simulations

MD simulations were performed using GROMACS2022 [19] utilizing AMBER ff19SB [20] and GAFF2 [21] force field parameters for proteins and ligand, respectively. The force field parameters for the chromophore were generated using the antechamber tool of AmberTools22. Bonded and short-range nonbonded interactions were calculated every 2 fs, and periodic boundary conditions were employed in all three dimensions. The particle mesh Ewald (PME) method [22] was used to calculate long-range electrostatic interactions with a grid density of 0.1 nm^−3^. A force-based smoothing function was employed for pairwise nonbonded interactions at a distance of 1 nm with a cutoff of 1.2 nm. Pairs of atoms whose interactions were evaluated were searched and updated every 20 steps. A cutoff (1.2 nm) slightly longer than the nonbonded cutoff was applied to search for the interacting atom pairs. Constant pressure was maintained at a target of 1 atm using the Parrinello-Rahman algorithm [23]. Temperature coupling was maintained at 300 K with the V-rescale algorithm [24]. The iNicSnFR12 system was generated by computationally mutating E476K, N11E, R467S, Q431D in iNicSnFR3a.

### Dynamic network analysis

Dynamic network analysis for iNicSnFR3a and iNicSnFR12 in the nicotine-bound form was performed using the NetworkView plugin in VMD [25–27]. For each system, a network map or graph was created where nodes are the Cα atoms of each protein residue. The edges are added between a pair of nodes if the heavy atoms of the protein residues assigned to these nodes are within 4.5 Å distance for > 75% of the simulation time. The edge weights *w_ij_* are derived from pairwise correlations (*C_ij_*) which define the probability of information transfer across a given edge:

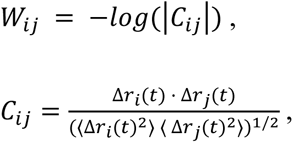

where Δ*r_i_*(*t*) = *r_i_*(*t*) – <Δ*r_i_*(*t*)> and *r_i_*(*t*) is the position of the atom corresponding to the *i*^th^ node. The resulting network was used to determine the optimal path between two given nodes (a source and a sink), using The Floyd–Warshall algorithm [https://doi.org/10.1145/367766.368168]. The optimal path is defined to be the connecting route between the two nodes (residues), which minimizes the number of intermediate nodes and maximizes the sum of edge betweenness of the connecting route.

### Differential scanning fluorimetry and intrinsic fluorescence melts

Differential scanning fluorimetry (DSF) [28] was used to evaluate the thermal stability of iNicSnFR variants. In the DSF assay, samples containing 45 μL of 5 μM purified protein (3×PBS, pH 7.0) with and without nicotine at twice the respective variant’s EC_50_ value were mixed with 5 μL of a stock solution of 200×SYPRO^®^ Orange dye (Invitrogen, Eugene, OR) in triplicate. Blank samples with the same concentrations of nicotine and SYPRO^®^ Orange dye were also prepared in triplicate for blank subtraction. All samples were assayed on a CFX96 Real-Time PCR Detection System (Bio-Rad Laboratories, Hercules, CA). Samples underwent a thermal melt from 25 °C to 99 °C with temperature holds every 30 s at 0.5 °C increments. Fluorescence values were measured in relative fluorescence units (RFU) using the FRET scan mode. Backgrounds were subtracted from all samples.

Since the iNicSnFR variants have intrinsic fluorescence arising from the cpGFP moiety which overlaps with that of SYPRO^®^ Orange dye, intrinsic fluorescence melts were performed in triplicate and the average fluorescence values were subtracted from the DSF samples. For the intrinsic fluorescence samples, 5 μL of buffer (3×PBS, pH 7.0) was added to the protein samples in place of SYPRO^®^ Orange dye. The intrinsic fluorescence samples underwent the same thermal melts as the DSF samples, and blank subtraction was performed. Apparent melting temperatures were determined from peaks in the first derivative plots of the melting curves that had average intrinsic fluorescence subtracted out (see Figures S3, S4).

## Results

### Computational prediction

During the development of iNicSnFR3, the ligand binding pocket between the two lobes of the PBP Venus flytrap domain was previously adapted to nicotine by site-saturated mutagenesis (SSM) on “first-shell” residues that lie within 7 Å of the bound ligand as well as on several “second-shell” residues that interact with the first-shell residues [1, 6]. Because (a) mutations at greater distances from the binding site sometimes affect ligand binding and/or enzymatic activity [29, 30], and (b) mutations computationally designed for enhanced stability often show enhanced function [31], we applied computational design to identify possible mutation hotspots outside the first and second shell.

We first used AutoDock Vina to dock nicotine into iNicSnFR3a [32]. The structure of iNicSnFR3a was extracted from the crystal structure of varenicline-bound iNicSnFR3a (PDB ID: 7S7T) and the varenicline molecule was deleted from the structure to provide an empty binding pocket for nicotine docking. The structure was prepared in AutoDockTools by removing waters, adding polar hydrogens, and assigning Gasteiger charges. To verify the docking procedure, we first docked varenicline into the prepared structure, where the highest scoring pose showed ∼ 0.5 Å RMSD from the original varenicline molecule in 7S7T. Then, we docked nicotine into the structure and the conformation with the highest score was selected for computational design.

The nicotine-docked iNicSnFR3a structure was first standardized by the computational protein design suite TRIAD. Then we used the single mutation stability module from TRIAD to rank every possible amino acid substitution at every residue position in the PBP moiety of iNicSnFR3a, one at a time, by predicting the change in energy of folding upon mutation. The top 100 variants were selected (**Table S2)**, from which we selected 34 positions for subsequent sequence design: S9, N20, E24, E27, R36, N39, N46, K51, R52, E64, E78, S325, K336, R341, K342, K362, L384, T413, K414, M418, E429, Q431, D434, D439, D452, D453, K454, R467, E476, N497, K499, D501, E513, E517, excluding all prolines and positions that have been optimized in previous evolution [1].

Finally, we applied the sequence design module from TRIAD to these selected positions individually. All 20 amino acid residues were simulated at each position and the resulting energy changes were evaluated. This is essentially the same prediction strategy as used in the TRIAD single mutation stability module, but the overall energy was calculated more comprehensively and reliably using the Rosetta algorithm. Sequence design on each selected position generated 20 entries of energy scores, from which we identified the positions that could generate the largest energy change when mutated from their wild-type sequences (termed “energy gain”, **Table S2)**. The top 10 positions (N20, E27, N46, K51, R52, K342, Q431, K454, R467, and E476), which gave the largest energy changes from wild-type sequence upon mutation, plus N11 which was mutated from iNicSnFR3a to iNicSnFR3b, were selected for SSM screening.

### Directed evolution

The evolution path from iNicSnFR3a [1] to iNicSnFR12 is shown in **Figure 1A**. In this study, we performed the first round of SSM on positions N20, K342, Q431, and R467 on iNicSnFR3a individually; but the mutations (N20R, K342Y, Q431A, and R467S) displayed little to no improved sensitivity. Therefore, we tested various combinations of these mutations via SDM. The double mutant (iNicSnFR3a Q431A R467S) showed slightly improved sensitivity.

**Figure 1.**
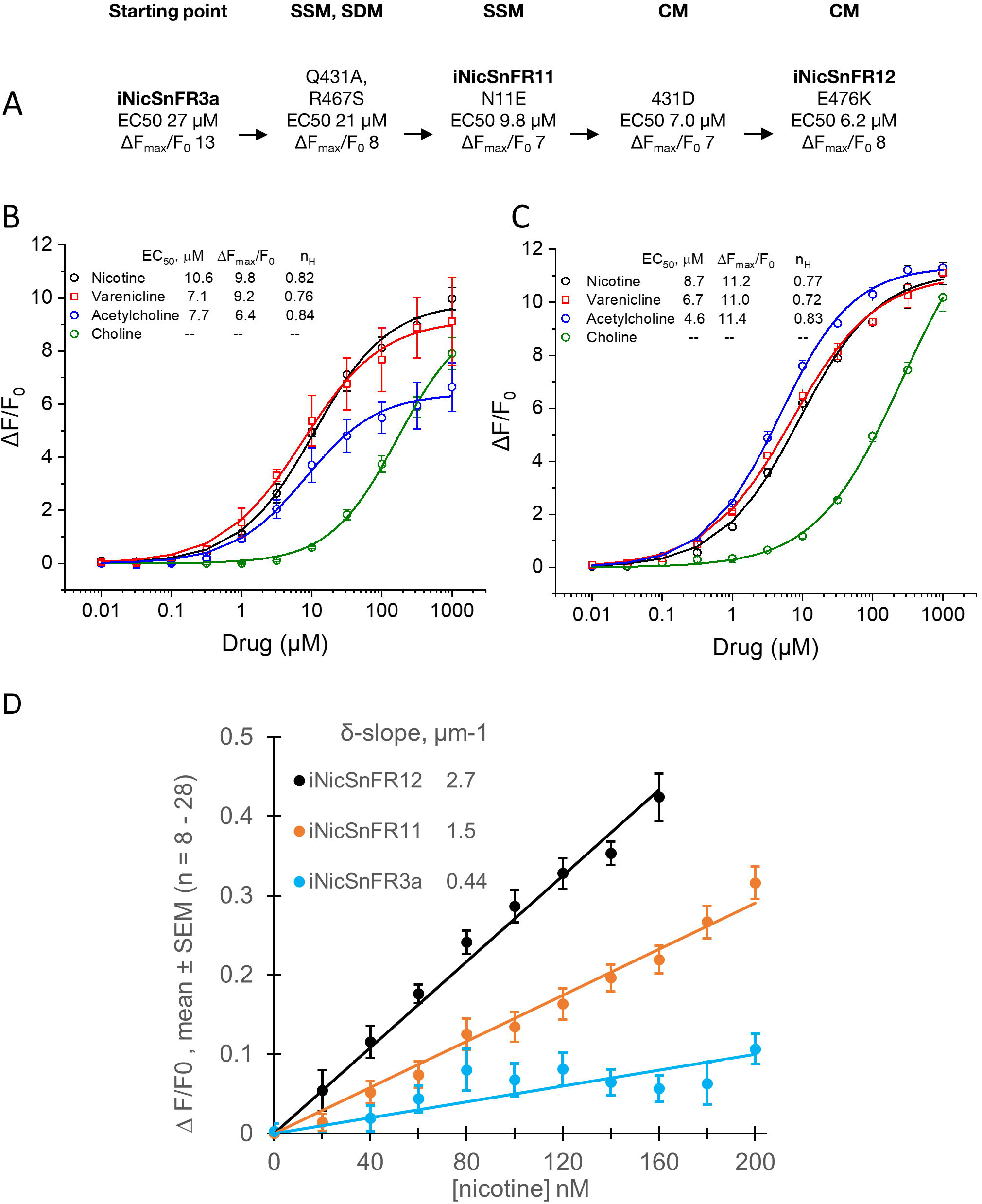
Directed evolution and fluorescence characteristics of iNicSnFR11 and iNicSnFR12. A. The path of protein engineering of iNicSnFR constructs. Data for iNicSnFR3a are the average of two measurements reported previously for the purified protein [1]. Other measurements reported in this figure were made on bacterial lysates, which often have different ΔF_max_/F_0_ and EC_50_ values than the purified proteins (compare with panels B and C). SSM, site saturation mutagenesis; SDM, site-directed mutagenesis; CM, combinatorial mutagenesis. B, **Dose-response relations for purified iNicSnFR11.** B1, Dose-response relations for several nicotinic drugs. Purified iNicSnFR11 was measured at 100 nM in 3×PBS, pH 7.40. Data are mean ± SEM (n = 3). **C. Dose-response relations for purified iNicSnFR12.** C1, Dose-response relations for several nicotinic drugs. Purified iNicSnFR12 was measured at 100 nM in 3×PBS, pH 7.40. Data are mean ± SEM (n = 3). **D, Dose-response relations for iNicSnFR variants at sub-μM [nicotine]**. Purified iNicSnFR3a, iNicSnFR11, and iNicSnFR12 were measured at 100 nM in 3×PBS, pH 7.40. The legend shows values for the fitted δ-slopes. Data points are mean ± SEM (n = 8 to 28).

Starting from iNicSnFR3a Q431A R467S, we performed the second round of SSM on N20 and K342 again, as well as on other positions from the list including N11, E27, N46, K51, R52, K454, E476. We identified N11E as a beneficial mutation, yielding iNicSnFR3a N11E Q431A R467S, which we termed iNicSnFR11.

However, the third round of SSM on iNicSnFR11 yielded no beneficial mutations. We reasoned that iNicSnFR11-nicotine binding affinity might be trapped at a local maximum [33]. We therefore constructed and screened variants that combined several of the previously identified beneficial mutations. We made the first such “combinatorial library” of iNicSnFR11 incorporating some or all of the following mutations: K342, Y342, Qδ431, A431, G431, D431, E431, R467, G467, S467, H467. Screening this library yielded iNicSnFR11 A431D with improved nicotine sensitivity. Then we made the second combinatorial library of iNicSnFR11 A431D incorporating K51, L51, Q51, R51, R52, V52, T52, E476, K476, M476, V476. We obtained iNicSnFR11 A431D E476K with further improved nicotine sensitivity, which we termed iNicSnFR12. Therefore. iNicSnFR12 is iNicSnFR3a with mutations: N11E, Q431D, R467S, and E476K.

### Fluorescence characteristics of iNicSnFR11 and iNicSnFR12

We used excitation at 485 nm, performed emission measurements at 510 nm, and generated dose-response relations for purified iNicSnFR11 and iNicSnFR12 against four nicotinic agonists: nicotine, ACh, choline, and varenicline. For nicotine at iNicSnFR11, we found an EC_50_ of 10.6 μM (Figure 1, B1), or a 2.5-fold improvement over iNicSnFR3a [1]. The iNicSnFR11 data show roughly equal EC_50_ values for varenicline, ACh, and nicotine (Figure 1, B1), which was also true for iNicSnFR3a [1].

In contrast, the EC_50_ for choline is > 17-fold greater than for nicotine (the fitted value of 170 μM should be considered approximate because the fluorescence did not approach a maximum at 1 mM, the highest value tested). Because we intend to apply iNicSnFR11 and iNicSnFR12 in the sub-μM range, we emphasize measurements in this range. As expected, the dose-response relation of iNicSnFR11, ΔF/F_0_ vs [nicotine], was linear in this range. We term the proportionality constant the δ-slope to emphasize that it is measured at low [nicotine] (< 0.1x EC_50_). We consider the measured δ-slope more relevant than the metric called the S-slope = (ΔF_max_/F_0_) / EC_50_. The ΔF_max_/F_0_ and EC_50_ values, which are fitted to the Hill equation from measurements over the entire dose-response relation, could be influenced by complexities at high [nicotine]. For iNicSnFR11, the δ-slope was 1.5 μM^-1^ (Figure 2D), representing a 3.4-fold improvement over the value measured for iNicSnFR3a, 0.44 μM^-1^.

**Figure 2.**
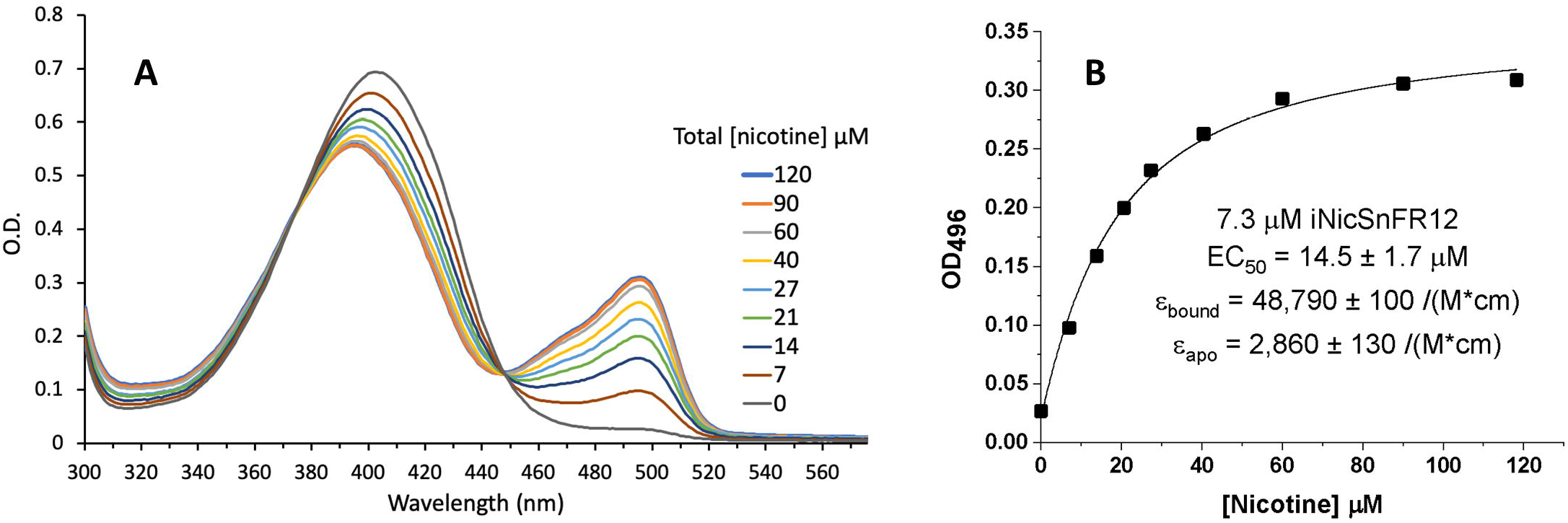
iNicSnFR12 shows increased absorption at 450-520 nm upon nicotine binding. A, Nicotine tartrate solution (1 mM) was added to the iNicSnFR solution (7.3 μM) to produce the indicated nominal [nicotine]. B, Fitting the OD at 496 nm (the local peak absorption) to a single binding model yielded the parameters indicated.

For iNicSnFR12, the EC_50_ for nicotine was 8.7 μM, the lowest value we have measured in the iNicSnFR series (**Figure 1, C1**). Varenicline and ACh had EC_50_ values similar (within two-fold) to the value for nicotine (**Figure 1, C1**); this similarity resembles the pattern for iNicSnFR3a and iNicSnFR11. Again, EC_50_ for choline was > 12-fold greater than for nicotine (because the dose-response relation for choline did not approach saturation at the highest [choline] tested, we provide no fitted EC_50_).

As expected, in the sub-μM range, nicotine displays a linear dose-response relation at iNicSnFR12 (Figure 1D). The δ-slope for nicotine is 2.7 μM^-1^. Therefore, iNicSnFR12 responds to nicotine with sensitivity representing a 6.1-fold improvement over iNicSnFR3a.

### Absorbance characteristics of iNicSnFR12

Figure 2A presents optical absorbance data for iNicSnF12 in the presence of various nominal [nicotine] between zero and the nearly saturating value of 120 μM at pH 7.0. Typical fluorescence experiments are conducted at excitation wavelengths between 470 and 500 nm. At the absorption peak (496 nM), the fitted molar absorptivity of the apo and nicotine-bound iNicSnFR12 are ε_apo_ = 2,900 M^-1^ cm^-1^ and ε_bound_ = 48,800 M^-1^ cm^-1^, respectively (Figure 2B). Because of uncertainties in concentrations, dilutions, and blanks, we consider that the ratio ε_bound_ / ε_apo_ is in the range of 14 to 21. The fitted K_d_ value is 14.5 μM (Figure 2B). In nicotine-induced fluorescence experiments on the same sample of purified iNicSnFR12 (diluted to 70 – 200 nM), we measured ΔF_max_ / F_0_ = 10.2 ± .8 and EC_50_ = 10.6 ± 1.2 μM (mean ± SEM, n = 3). The modestly greater (∼1.5-fold) EC_50_ value at pH 7.0 vs 7.4 is consistent with previous observations on increases in EC_50_ with lower pH for other iDrugSnFRs [1, 5] and may be explained by the role of protonation in the ”candle snuffer” mechanism [6].

### Fluorescence quantum yield of iNicSnFR3a and iNicSnFR12

The Φ for iNicSnFR3a (corrected as described in Methods) is 0.733; for iNicSnFR12, 0.783. We consider that these estimates are reliable within ±15%, so that Φ for iNicSnr3a and iNicSnFR12 are between 0.623 and 0.843 and between 0.666 and 0.901, respectively. Therefore, the Φ of the liganded iNicSnFR variants resembles that of EGFP itself (0.6) and may be slightly higher.

### Modeling of nicotine within the diffuse electron density

In our previous study [6], the X-ray structure of iDrugSnFR3a saturated with nicotine (PDB ID: 7S7U) was determined with diffuse electron density for the nicotine, which could not be definitively built in the deposited model. To better understand the structural basis for nicotine binding and selectivity of this structure, we sought a more precise ligand-bound model. We developed a computational workflow that involves 1) exhaustive exploration of drug conformations, 2) clustering of the energy minimized conformations, 3) launching of parallel MD simulations from each cluster, 4) re-clustering the MD data, and 5) analysis of cluster populations and cross-correlation calculations with the X-ray map (details in Methods) (Figure 4A). In our analysis, the most highly populated cluster (cluster 1) showed the greatest cross-correlations with the experimental map. Therefore, we chose the pose closest to the centroid of cluster 1 (pose 1) as the representative ligand-bound model, for further simulations and analysis. The existence of multiple sparsely-populated clusters might arise because nicotine is inherently dynamic in the binding pocket, a factor that might contribute to the diffuse electron density.

Next, we analyzed detailed interactions of nicotine within the binding pocket using all the members of cluster 1 (Figure 3B). The protonated amine at the methylpyrrolidine of nicotine formed several cation-π interactions with Y65, Y357 and Y460, and hydrogen bond interactions with the sidechain of D397 and the backbone carbonyl of N355. Additional π - π like interactions occurred between the pyridine group and F12, F391 and W436.

**Figure 3:**
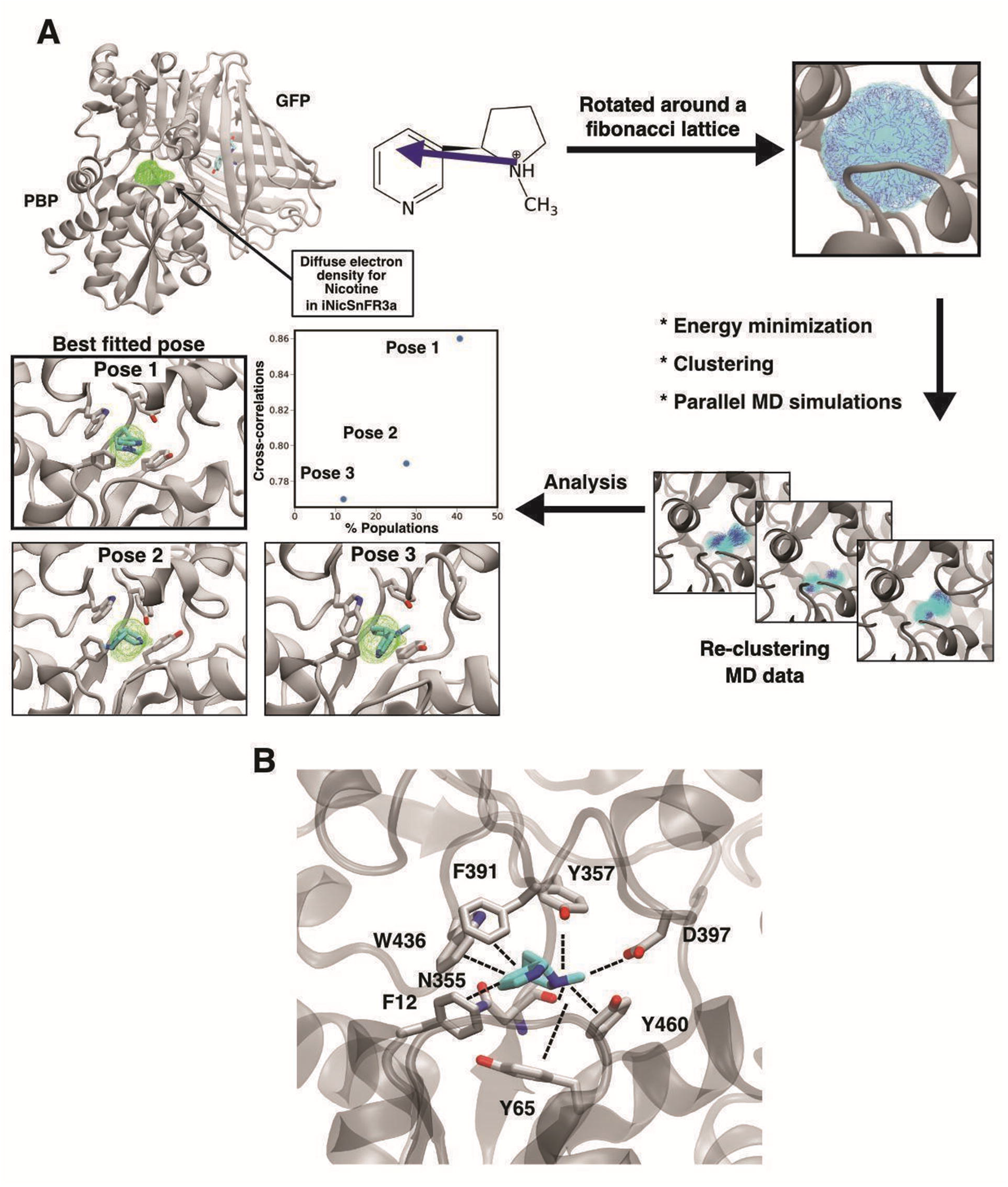
An MD simulation-based workflow to assign a nicotine-bound pose in the X-ray map of nicotine-bound iNicSnFR3a (PDB: 7S7U). (A) The PBP and the GFP (chromophore shown as sticks). The diffuse X-ray density of nicotine is shown in green. To search for possible nicotine poses in the binding pocket, a vector within the ligand was defined and the vector was rotated along a Fibonacci lattice, including both internal conformations and self-rotation. All the poses were then energy minimized, followed by clustering, and parallel MD simulations. The simulation data set was then re-clustered. The molecular view of the top 3 poses (defined by the member, closest to the centroid of a particular cluster) is shown. Pose 1, a member of the most highly populated cluster, overlaps best with the X-ray map as quantified by the map to model cross-correlation calculations. (B) Molecular view of the detailed ligand-protein interaction in Pose 1. A contact was considered to be present if any two atoms from nicotine and protein lie within 3.5 Å in > 30% of the total members of cluster 1.

**Figure 4.**
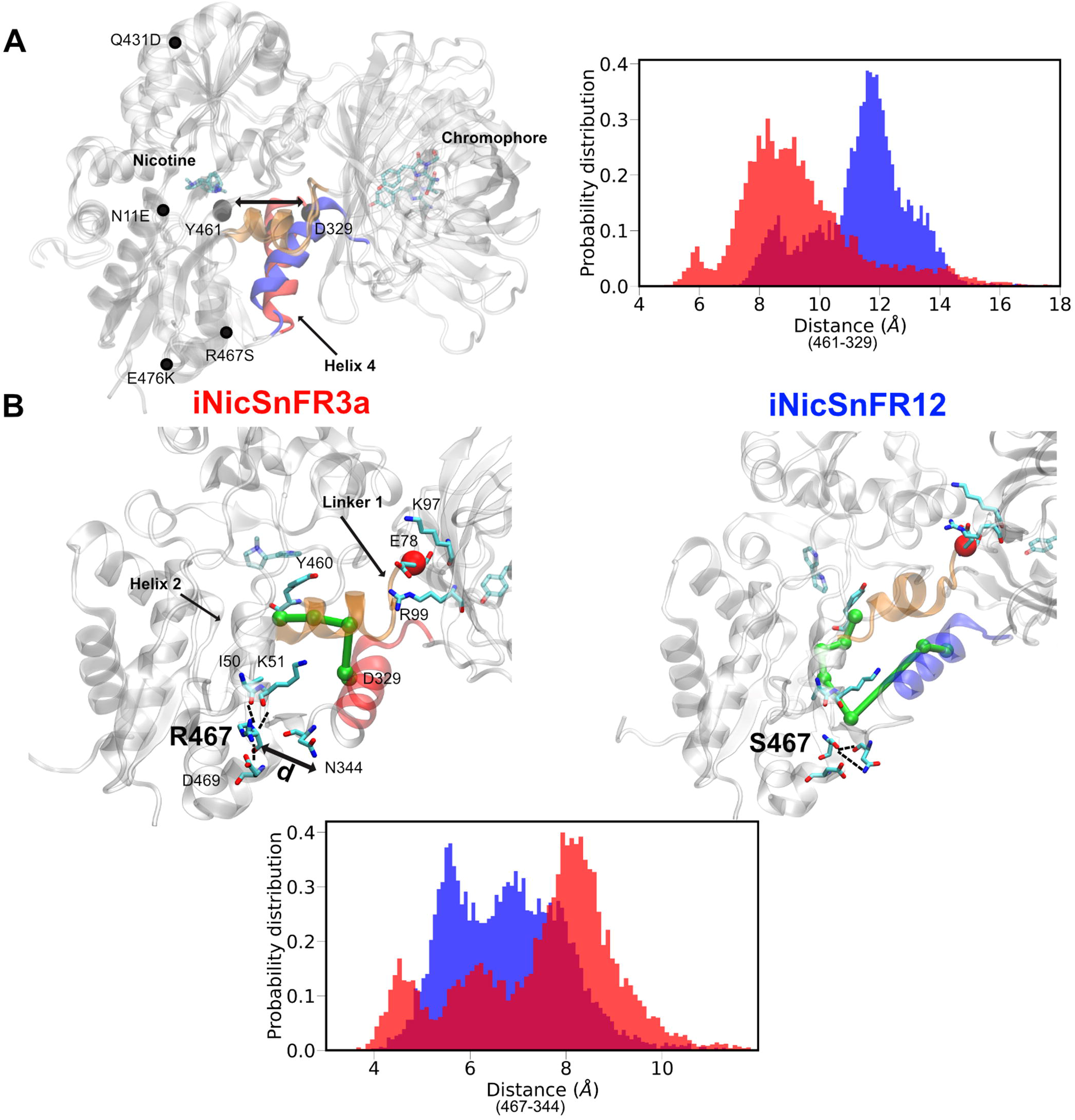
Tilting of linker helix 4 in iNicSnFR12 might modulate ligand binding. (A) All the mutations present in iNicSnFR12 from iNicSnFR3a are labeled. In iNicSnFR12, helix 4 (colored in blue) shifts towards the GFP. To measure this process, we analyzed the probability distribution of the distance between the Cα atom of a terminal residue D329 and the Cα atom Y461, the terminal of a β-sheet that locates nearest to helix 4, residue Y461. (B) The allosteric network from the nicotine binding site residue, Y460, to the N-terminal residue of helix 4, D329, is depicted in green. The probability distribution shows hydrogen bond interactions of residue 467 and N344, measured by calculating the distance between the side chains of these two residues in the two systems.

### Nicotine binding and structural changes

To visualize dynamic nicotine binding in both iNicSnFR12 and iNiCSnFR3a, we performed twelve replicate MD simulations for the two sensors, each for 1 μs, totaling 12 μs for each system, using the ligand-bound structure determined above. In the majority of replicates (8 out of 12 for each system), nicotine remained bound, indicating that the starting structure represented a comparably stable bound state for both sensors. We analyzed these replicates further to understand structural consequences of the differentiating mutations. The N-terminus of the TDPEGAYETVKKEYKR sequence comprises helix 4 in the nomenclature of Fukami-Kobayashi et al. [34, 35] and connects the PBP to GFP. Interestingly, in NicSNFr12, helix 4 was shifted ∼4 Å away from the centroid of the PBP (Figure 4A), compared to NicSNFr3a. One measure of this shift is the probability distribution of the distance between the Cα atom of a terminal residue D329 and the Cα atom Y461, the terminal of a β-sheet that locates nearest to helix 4, residue Y461 (Figure 4A). Next, we sought to investigate 1) the basis for this striking change, and 2) whether this shift could be related to increased fluorescence response to nicotine in iNicSnFR12.

Four mutations distinguish iNicSnF12 from the earlier iNicSnFR3a. Of these, R467S is located nearest to helix 4 and exhibited a differential hydrogen bond pattern in the two sensors (Figure 4B). In iNicSnFR3a, R467 reaches out to helix 2 located just “above” it and forms a helix capping interaction with the backbone of the two C-terminal residues I50 and K51 (in addition to a salt-bridge interaction with D469 which locates just “below” R467). These interactions are, however, absent in iNicSnFR12, which affects the thermal stability of the variant as described below. Instead, S467 interacts with the side-chain and backbone of helix 4 C-terminal residues N344 and W343 of helix 4, respectively. We quantified this observation by measuring the distance from the sidechain of residue 467 to N344 (Figure 4B). The probability distribution of this distance metric clearly showed a closer contact between S467-N344 in iNicSnFR12 compared to in iNicSnFR3a. This interaction slightly unwinds and pulls the C-terminus of helix 4 towards the centroid of the PBP **(Figure S2)**; this action could help to force the other end of the helix to tilt away from the PBP as a balancing effect.

We next focused our analysis on the hypothesis that the helix shift modifies sensitivity to nicotine in iNicSnFR12 by remodeling allosteric connectivity to or from the ligand binding site. We presume this connectivity can be partially evaluated in terms of the pathways of residues that mediate force propagation among functional sites in the protein [25]. To compare these pathways, we performed dynamic network analysis on both the iNicSnFR3a and iNicSnFR12 systems using the Network-View plugin in VMD (see Methods) [25–27]. The source was selected as Y460, a key aromatic residue coordinating the cationic amine of nicotine as described above (Figure 3B). The sink was selected as D329, located at the N-terminal tip of the tilted helix 4. Our analysis showed a dramatic change in the allosteric pathways between these two residues. In iNicSnFR3a, the pathway traversed the short helix adjacent to linker 1 (orange). Linker 1 connects the PBP to the GFP and includes the critical residue E78. In the apo state, E78 lies within ∼ 2.5 Å of the fluorophore, acting as a “candle snuffer” that drastically decreases fluorescence. The conformational change upon ligand binding moves E78 ∼ 14 Å away from the fluorophore, allowing fluorescence [6]. In the bound state, E78 forms salt bridges with Lys97 and Arg99 [6]. In the MD simulations of the iNicSnFR12 bound state, these salt bridges remained present; and the “candle snuffer” mechanism therefore remained intact. However, the mechanical pathway between Y460 and D329 traversed the PBP strand terminating in the mutated position R467S.

### Thermal stability characterization of iNicSnFR3a, iNicSnFR12, and iNicSnFR12-S467R

To explore the stability implications of the R467S mutation in iNicSnFR12 that putatively deletes the helix capping interaction between the sidechain of R467 and the C-terminal residues of helix 2, differential scanning fluorimetry (DSF) was performed on samples of iNicSnFR3a, iNicSnFR12, and iNicSnFR12-S467R at pH 7.0 in the absence of ligand (Figure 5A). iNicSnFR12-S467R differs from iNicSnFR12 by a single S467R point mutation, which reintroduces the putative helix capping interaction. The three variants exhibited two distinct melting transitions (see **Figures S3, S4**). The unfolding of the PBP domain occurred at the lower apparent melting temperature, T_m_(low), while the unfolding of the GFP moiety occurred at the higher apparent melting temperature, T_m_(high) (**Figures S3, S4**). While the T_m_(high) values for iNicSnFR3a, iNicSnFR12, and iNicSnFR12-S467R were indistinguishable, the T_m_(low) value associated with PBP unfolding was lower for iNicSnFR12 compared to iNicSnFR3a and iNicSnFR12-S467R (Figure 5A). This observation supports the hypothesis that R467 plays an important role in stability of the PBP. To determine the effect of nicotine on the stability of these variants, DSF was performed on each variant with a concentration of nicotine equal to twice the respective EC_50_ values (Figure 5B). In nicotine-induced fluorescence experiments on the sample of purified iNicSnFR12-S467R, we measured an EC_50_ of 25.6 ± 1.8 μM (mean ± SEM, n = 3). The presence of nicotine at twice the EC_50_ concentrations increased values for T_m_(low) ranging from 1.7 °C to 3.3 °C and decreased values for T_m_(high) by less than 1 °C. Nicotine, therefore, has an overall stabilizing effect on the PBP domains of the tested iNicSnFR variants.

**Figure 5.**
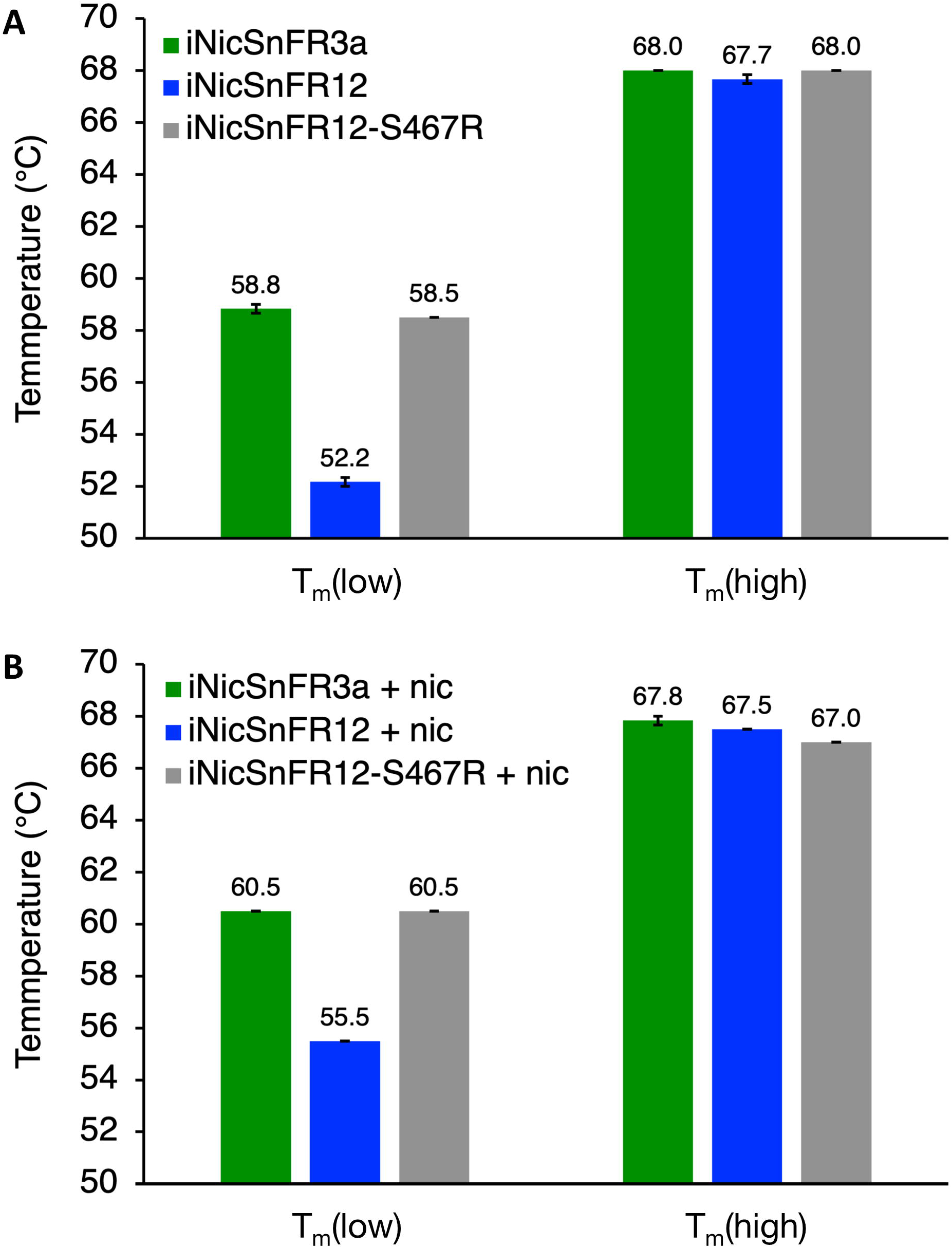
S467R mutation to iNicSnFR12 restores enhanced stability inherent to iNicSnFR3a. A, DSF for iNicSnFR3a, iNicSnFR12, and iNicSnFR12-S467R in 3×PBS, pH 7.0 in the absence of ligand. Tm(low) values were determined to be 58.8 ± 0.2 °C, 52.2 ± 0.2 °C, and 58.5 ± 0.0 °C for iNicSnFR3a, iNicSnFR12, and iNicSnFR12-S467R respectively. Tm(high) values were determined to be 68.0 ± 0.0 °C, 67.7 ± 0.2 °C, and 68.0 ± 0.0 °C for iNicSnFR3a, iNicSnFR12, and iNicSnFR12-S467R respectively. B, DSF for iNicSnFR3a, iNicSnFR12, and iNicSnFR12-S467R at pH 7.0 with [nicotine] = 2×EC_50_. Tm(low) values were determined to be 60.5 ± 0.0 °C, 55.5 ± 0.0 °C, and 60.5 ± 0.0 °C for iNicSnFR3a, iNicSnFR12, and iNicSnFR12-S467R respectively. Tm(high) values were determined to be 67.8 ± 0.2 °C, 67.5 ± 0.0 °C, and 67.0 ± 0.0 °C for iNicSnFR3a, iNicSnFR12, and iNicSnFR12-S467R respectively. Data are mean ± SEM, n = 3.

### iNicSnFR12 detects nicotine at 100 nM concentrations in rat and human serum

Figure 1, **C2** shows that solutions containing iNicSnFR12 detect nicotine at < 100 nM. We tested whether this capability also applies to solutions containing rat or human biofluids. Firstly, in PBS alone, fluorescence increased as the percentage of rat serum increased, possibly due to the presence of bilirubin. Secondly, we define F_0_’ as the fluorescence in PBS plus rat serum plus iNicSnFR12. F_0_’ was greater than the arithmetic sum of F_0_, plus the fluorescence of 10% rat serum alone, by a factor 3.52 ± 0.1. This suggested that rat serum contains endogenous molecule(s) that activate iNicSnFR12 in the absence of nicotine. With heat-treated serum, the ratio was only slightly less (3.22 fold). This suggested that the elevated F_0_’ arises from a small molecule(s) in serum. We infer that the endogenous non-protein ligand(s) in rat serum activate iNicSnFR12 to 31% (3.52/11.2; see Figure 2, **C1**) of the maximum for nicotine.

Thirdly, despite the elevated F_0_’, in PBS containing 10% rat serum, iNicSnFR12 displayed fluorescence responses at nicotine concentrations as low as 50 nM nicotine (Figure 6A). The small variability and linearity suggest that the limit of detection is somewhat lower, perhaps 20 nM. Under these conditions, the δ-slope had a deceased value, 0.37 ± 0.02 µM^-1^; we term this parameter the δ’-slope, and it is several fold lower than the zero-serum δ-slope value of 2.7 μM^-1^ (Figure 1D).

**Figure 6.**
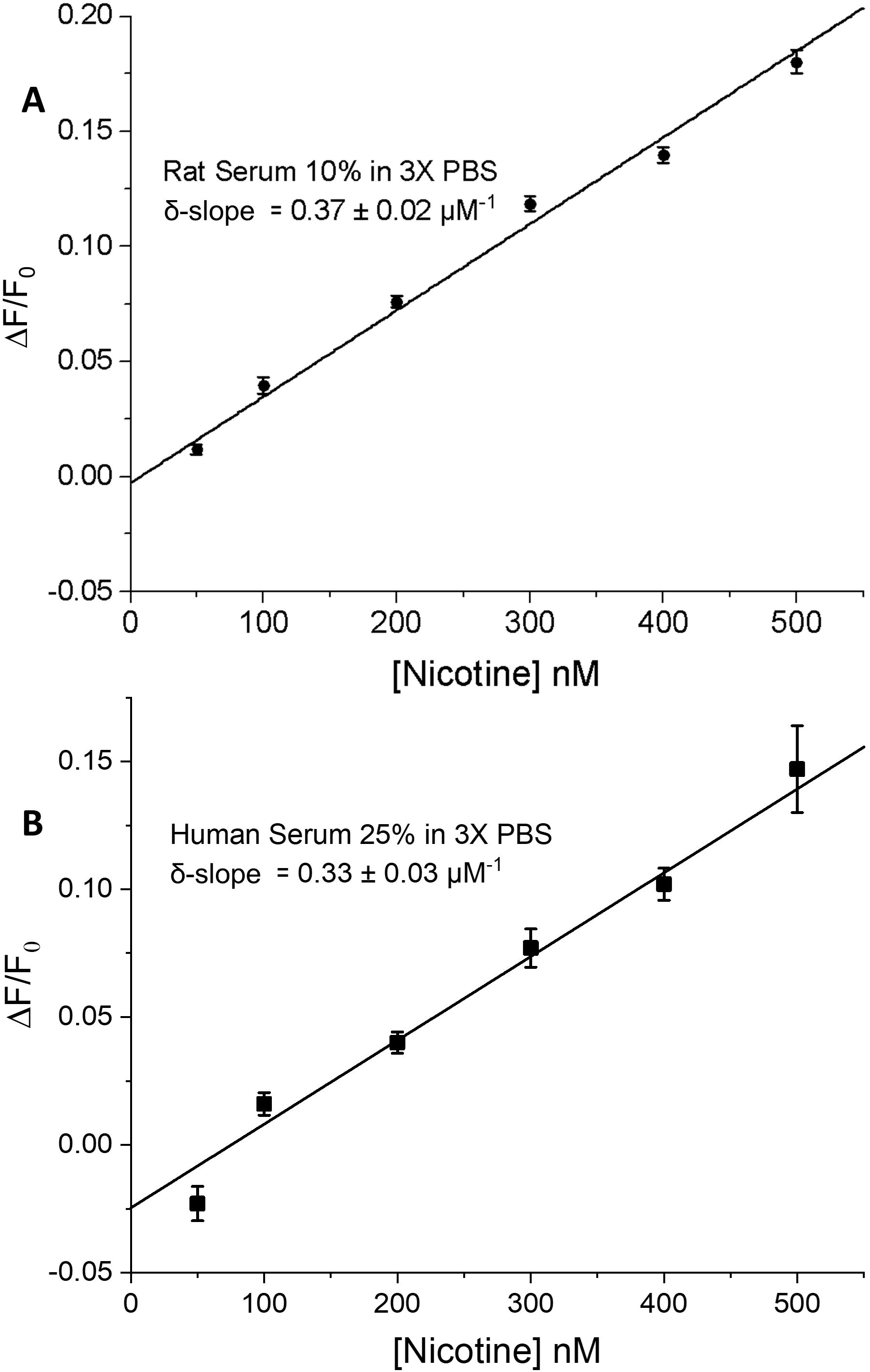
Dose-response data for nicotine at iNicSnFR12 in biofluids. **A**, Rat serum, 10% in 3X PBS. iNicSnFR concentration was 100 nM. Data were measured 5 - 30 min after addition of nicotine. The fitted line corresponds to δ’-slope = 0.52 ± 0.01 μM^-1^. Data are mean ± SEM, N = 9 samples per concentration. **B**, Human serum, 25% in 3X PBS. iNicSnFR concentration was 100 nM. Data were measured 15 - 30 min after addition of nicotine. The fitted line corresponds to δ’-slope = 0.33. ± 0.03 μM^-1^. Data are mean ± SEM, N = 9 samples per concentration.

We simulated the fluorescence responses in the presence of an endogenous ligand. The relationship between δ -slope and decreased δ’-slope was consistent with simulations of partial activation by endogenous ligand(s) (**Figure S5**).

PBS containing 25% human serum from nonsmokers had an intrinsic fluorescence 2.0 ± 0.1 fold larger than PBS alone. In 25% human serum plus iNicSnFR12, F’_0_ was 1.9 ± fold higher than the arithmetic sum of F_0_ in PBS and the fluorescence of 25% human serum. This led us to define a new parameter, F_0_’, suggesting that human serum, like rat serum, has endogenous ligand(s) that activate iNicSnFR12. The calculated activation is 17% (1.9/11, see Figure 2,C1) of the maximum for nicotine.

We tested responses to nicotine at iNicSnFR12 in PBS containing 25% serum (Figure 6B). The data had greater variance than in 10% rat serum, possibly because of the elevated fluorescence of serum alone. The measurements at 50 nM nicotine had a negative ΔF/F_0_, distorting the extrapolated zero-concentration value. Therefore, in 25% human serum, the elevated F’_0_ and resulting variability lead to a limit of detection limit of ∼ 100 nM. The measured δ’-slope was 0.33 μM^-1^, decreased from the zero-serum value of 2.7 μM^-1^ (Figure 1D).

These preliminary data show that iNicSnFR12 responds to nicotine in two biofluids, with a reduced sensitivity (termed S’-slope), as expected from the elevated F_0_. See Discussion for the possible nature of the endogenous ligand(s).

## Discussion

### Characteristics of iNicSnFR12

We show an iterative approach to designing and testing improved fluorescent sensors for nicotine. The approach includes computational protein design and modest-throughput mutagenesis screening using iNicSnFR fluorescence to identify successful variants. We also show a systematic, internally consistent description of nicotine interaction with designed protein sensors, and molecular dynamics simulations to visualize structural consequences of sensitizing mutations on allosteric connectivity involving the ligand binding site.

Computational approaches in this work enabled the identification not only of potentially sensitizing mutations, but also of an experimental nicotine binding pose, and of the structural impact of mutations lacking structural data. These results, however, should be interpreted with appropriate caution, given that large-scale structural impact of engineered mutations may be insufficiently sampled on the timescale of an atomistic simulation, and mechanisms of activation cannot be conclusively determined in absence of experimental structures for both variants in multiple functional states. Classical molecular dynamics force fields may be particularly limited in modeling cation-π interactions, as they do not explicitly consider quantum mechanical terms [36, 37]. Still, substantial remodeling of allosteric pathways involving ligand binding is consistent with a dramatically altered conformational landscape for the nicotine-bound state of iNicSnFR12 versus iNicSnFR3a (Figure 4).

R467S is one of the mutations that distinguish iNicSnFR12 from previous iNicSnFR variants (Figure 1A). Comparative thermal stability data for Arg and Ser at position 467 support the idea that the side chain at position 467 affects the stability of Apo-iNicSnFR12 via the presence (Arg) or absence (Ser) of a helix capping interaction (Figure 5). Also, at least part of the increased nicotine sensitivity of iNicSnFR12 correlates with the increased thermal stability conferred by nicotine binding to the Ser-467 variant (Figure 5).

Our measurements represent the first data comparing the absorption characteristics of an iDrugSnFR between the apo and drug-bound state (Figure 2). The ε_apo_ and ε_bound_ values resemble those for various GCaMP constructs, which also have cpGFP moieties [10, 38]. For iNicSnFR12, the absorption ratio ε_bound_/ε_apo_ is in the range of 14 to 21, whereas the fluorescence data yield an F_max_/F_0_ ratio of 10.2. It should be noted that the absorption data were taken at iNicSnFR12 concentrations 37- to 90-fold higher than the fluorescence experiments, so that the ε_apo_ and F_0_ values may be separately distorted by systematic errors that affect these ratios. The good agreement between the EC_50_ from fluorescence measurements and the K_d_ from absorption experiments supports the idea that the same binding event and conformational change (movement of the candle snuffer [6]) governs both the fluorescence measurements and absorption changes, by perturbing the hydrogen bonds around the chromophore [38, 39]. In the 400-450 nM range, iNicSnFR12 shows only minor ligand-induced changes in absorption, like other cpGFP-based sensors [38].

We also present the first measurements of fluorescence quantum yield, Φ, on an iDrugSnFR. We find that Φ for nicotine-bound iNicSnFR3a and iNicSnFR12 is at least as great as that for EGFP itself and exceeds 0.6, implying that further attempts to improve the quantum yield would yield < 1.7-fold enhancements in signal. The rather low ε_apo_ values for the iNicSnFR family vitiate quantum yield measurements for the unbound state. For the present data, we cannot determine whether the absorption and fluorescence of the unbound state arise (a) completely from a small fraction of the protein molecules in the active conformation, versus (b) a small amount of absorption and fluorescence by the protein in the “snuffed” conformation. We do know that ΔF_0_ becomes larger at higher pH, presumably because deprotonation of Glu78 drives the apoprotein toward the active conformation [6], suggesting that mechanism (a) dominates.

### Contemplated uses for iNicSnF12

Nicotine sensors are selected/designed to bind nicotine; we have not selected against sensitivity to ACh or varenicline. The constructs do retain sensitivity to varenicline and acetylcholine. Neither of these molecules are expected to interfere with contemplated continuous measurements in human biofluids such as sweat and interstitial fluid.

Selection against choline sensitivity has been a secondary goal. The concentration of choline in healthy adult human and rat plasma is 10 μM [40–44]. The EC50 for choline at iNicSnFR12 is at least 100 μM (Figure 1**, C1**); therefore, [choline] in 10% rat serum and 25% human serum is 1μM and 2.5 μM, respectively. These [choline] are expected to produce at most 1% and 2.5% activation at iNicSnFR12, respectively. The measured activation (F_0_’/ F_0_) produced by the serum samples is > an order of magnitude higher (31% and 17% for 10% rat and 25% human serum, respectively.

These data suggest that another endogenous compound(s) activates iNicSnFR12 in rat or human serum. These compounds could be biogenic amines other than choline. In humans, the elevated F_0_ could also arise from the presence of varenicline. It remains to be tested whether the human biofluids of most interest, sweat and interstitial fluid, have similar concentrations of endogenous activated ligand(s). If so, identifying the endogenous ligand(s), and evolving iNicSnFR variants with decreased sensitivity to these ligands, remains a goal of further improvements in the iNicSnFR series.

A related goal is to selectively improve the sensitivity to nicotine so that ∼ 10 nM gives a larger signal. This would allow an iNicSnFR to become the sensor protein in minimally invasive or non-invasive devices that continuously monitor nicotine concentration in biofluids. The encouraging data with rat serum (Figure 6A) suggest that it would be worthwhile to measure [nicotine] continuously in rats performing intravenous nicotine self-administration [45].

In human pharmacokinetic data, smokers and vapers have as much as ten-fold variability in their nicotine half-life, in part because of polymorphisms in cytochrome P450 2A6 [46]. This variability also may inform the tactic that an individual uses to quit smoking, *e.g.* nicotine patch or varenicline [47]. A continuous nicotine monitor with sensitivity down to 10 nM would enable one to test the personal pharmacokinetics of nicotine during smoking or vaping sessions, as well as during the decay phase that continues for several hours after a session.

## Data availability

The data that support the findings of this study are available from the corresponding author upon reasonable request. The raw MD simulation trajectories can be found at 10.5281/zenodo.7529914

## Acknowledgements

A.L.N. was supported by California Tobacco-Related Disease Research Program (TRDRP) Grant 27FT-0022.

H.A.L. was supported by California TRDRP Grant 27IP-0057, National Institute on Drug Abuse Grant DA049140, and National Institute of General Medical Sciences (NIGMS) Grant GM-123582.

D.A.D. was supported by California TRDRP Grant T29IR0455.

N. J. F. was supported the Biotechnology Leadership Program through the Rosen Bioengineering Center at Caltech.

H. A. L. and S. M. were supported by a grant from the Merkin Institute for Translational Research at Caltech.

E. J. F. was supported by a Caltech Summer Undergraduate Research Fellowship from Paraskeva N. Danailov.

MD simulations were performed using the computing facilities of Karolina through EuroHPC (grant no. EHPC-REG-2021R0074) and Swedish National Infrastructure for Computing (SNIC2022/3-40), and supported by BioExcel.

N. H. was a supported by a Marie Sklodowska-Curie Postdoctoral Fellowship (grant no. 101107036).

R. J. H. and E. L. were supported by the Knut and Alice Wallenberg Foundation, the Swedish Research Council (2019-02433, 2021-05806), the Swedish e-Science Research Centre (SeRC), and the BioExcel Center of Excellence (EU-823830).

We thank Zoe Beatty, Kallol Bera, Heather Lukas, Anand Muthusamy, and Steven H. Zeisel for advice and guidance. We thank Purnima Deshpande for lab management.

## Responsible Editor

“If you are submitting a final version of a manuscript, please confirm that you have added the name of the PEDS Board Member responsible for editing it at the end of the manuscript.” Roberto Chica is the PEDS Editor / Associate Editor who signed the Decision Letter.

