## Supplementary Figures and Tables for "Interactive computational and experimental approaches improve the sensitivity of periplasmic binding protein-based nicotine biosensors for measurements in biofluids"

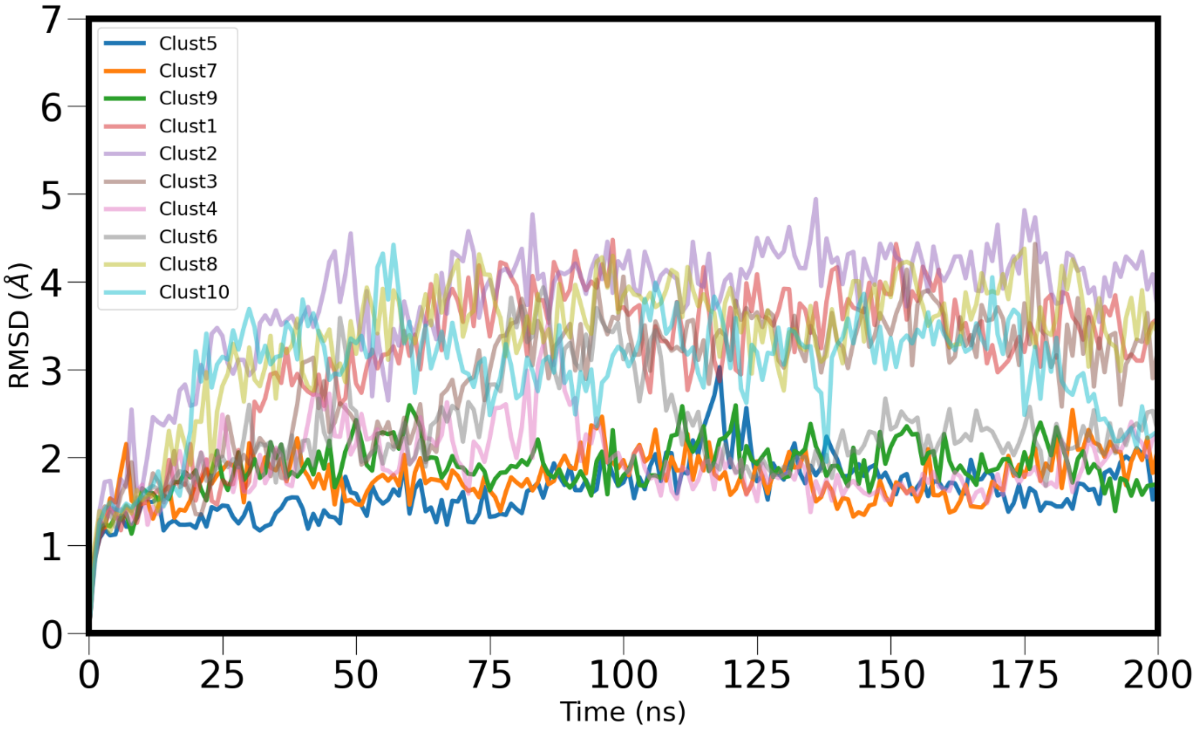


**Fig S1. Stability of PBP complexes in simulations.**

Root mean squared deviation (RMSD) of the PBP (after aligning the PBP to the initial frame) of the 10 nicotine-iDrugSnFR3a complexes, during the search for nicotine docking poses. Darker colors are used for the three trajectories that consistently stayed close to the holo conformation, while brighter curves illustrate simulations where the PBP binding cleft is (partially) opening towards the apo conformation.

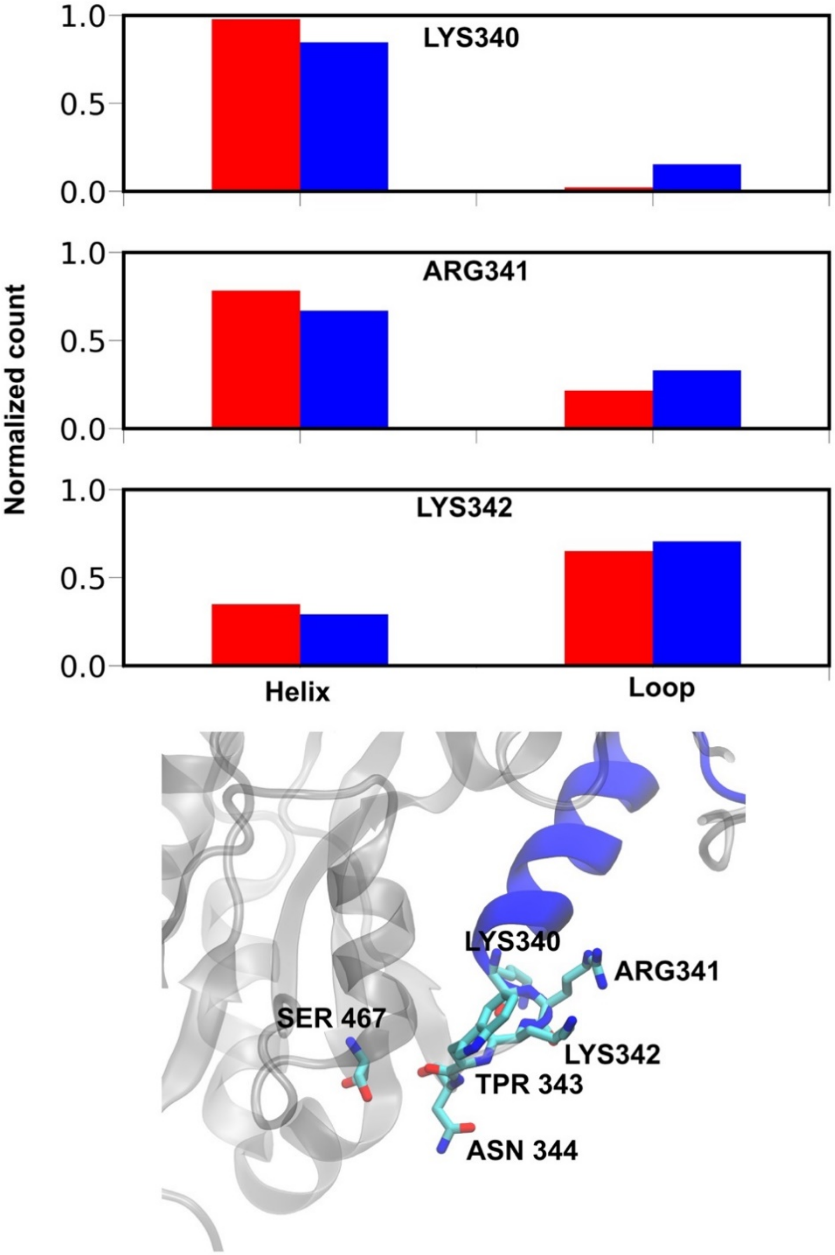


**Figure S2: Partial unwinding of Helix 4 in iNicSnFR12.**

Due to the interactions of SER467 with ASN344, located at the C-terminus of helix 4, this part of helix 4 spends a slightly lower fraction of the simulation in well-defined helical secondary structure in iNicSnFR12 (blue bars) in comparison to iNicSnFR3a (red bars), indicative of stretching/partial unwinding of the helix.

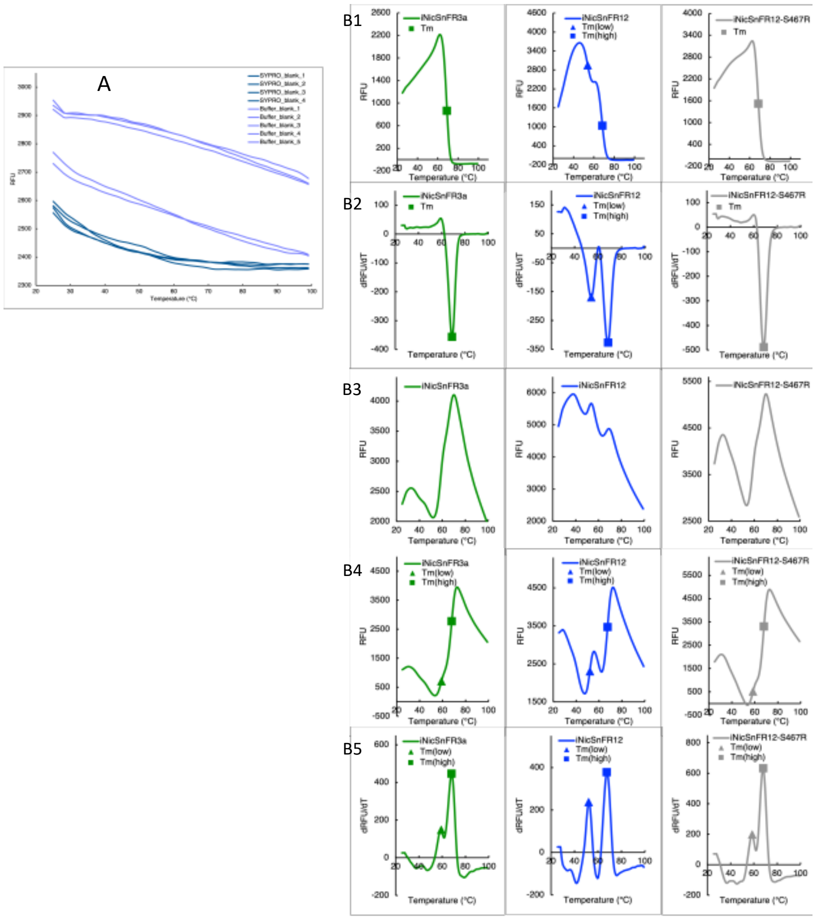


**Figure S3. Thermal melt data in the absence of nicotine.**

**A. Fluorescence of blank samples.**

Samples with the “SYPRO” and “Buffer” prefixes are with or without SYPRO® Orange, respectively.

**B1. Intrinsic fluorescence.**

Fluorescence values are averages of technical triplicates. Values for $T_m$, $T_m$(low), and $T_m$(high) are associated with decreases in intrinsic fluorescence due to GFP unfolding and were determined from the local minima of dRFU/dT curves for each replicate, the averages of which are presented in **Figure S3B2**. $T_m$ for iNicSnFR3a was 69.0 ± 0.0 °C. $T_m$(low) and $T_m$(high) for iNicSnFR12 were 54.0 ± 0.0 °C and 68.3 ± 0.2 °C, respectively. $T_m$ for iNicSnFR12-S467R was 68.5 ± 0.0 °C. Data are mean ± SEM, n = 3.

**B2. GFP apparent melting temperature determination.**

Values of dRFU/DT are averages of numerical derivatives taken on technical replicates.

**B3. Background-subtracted DSF.**

**B4. DSF minus intrinsic fluorescence.**

Average intrinsic fluorescence was subtracted from the fluorescence of triplicate DSF samples. Averages are shown. Values for $T_m$(low), and $T_m$(high) were determined from the peaks of dRFU/dT curves for each replicate, the averages of which are presented in **Figure S3B5**. Values for $T_m$(low) and $T_m$(high) are summarized in **Figure 5A**.

**B5. Apparent melting temperature determination.**

Values of dRFU/DT presented are averages of numerical derivatives taken on technical triplicates.

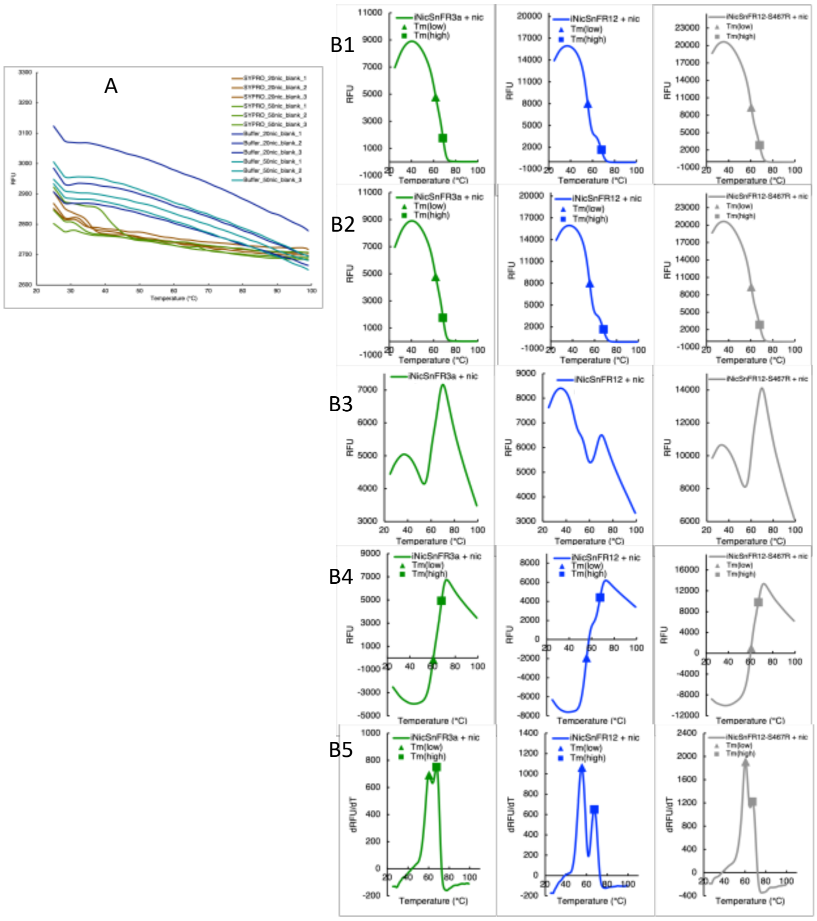


**Figure S4. Thermal melt data in the presence of nicotine.**

**A. Fluorescence of blank samples with nicotine.**

Samples with the "SYPRO" and "Buffer" prefixes are with or without SYPRO® Orange, respectively. Concentrations of nicotine are in µM (e.g., "20nic" refers to 20 µM nicotine).

**B1. Intrinsic fluorescence with nicotine.**

Fluorescence values are averages of technical triplicates. Values for $T_m$, $T_m$(low), and $T_m$(high) are associated with decreases in intrinsic fluorescence due to GFP unfolding and were determined from the local minima of dRFU/dT curves for each replicate, the averages of which are presented in **Figure S4B2**. $T_m$(low) and $T_m$(high) for iNicSnFR3a were 61.8 ± 0.2 °C and 68.5 ± 0.0 °C, respectively. $T_m$(low) and $T_m$(high) for iNicSnFR12 were 55.5 ± 0.0 °C and 68.0 ± 0.0 °C, respectively. $T_m$(low) and $T_m$(high) for iNicSnFR12-S467R were 60.3 ± 0.2 °C and 68.0 ± 0.3 °C, respectively. Data are mean ± SEM, n = 3.

**B2. GFP apparent melting temperature determination with nicotine.**

Values of dRFU/DT are averages of numerical derivatives taken on technical replicates.

**B3. Background-subtracted DSF with nicotine.**

**B4. DSF minus intrinsic fluorescence, each with nicotine.**

Average intrinsic fluorescence was subtracted from the fluorescence of triplicate DSF samples. Averages are shown. Values for $T_m$(low), and $T_m$(high) were determined from the peaks of dRFU/dT curves for each replicate, the averages of which are presented in **Figure S4B5**. Values for $T_m$(low) and $T_m$(high) are summarized in **Figure 5B**.

**B5. Apparent melting temperature determination with nicotine.**

Values of dRFU/DT presented are averages of numerical derivatives taken on technical triplicates.

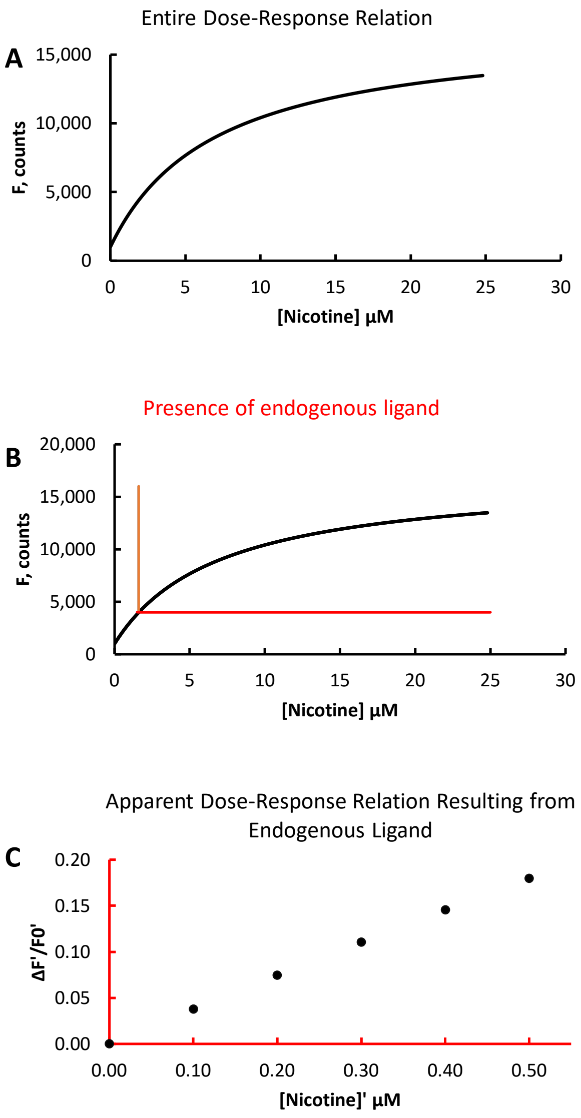


**Figure S5 Simulations of decreased S-slope caused by partial activation.**

We assume a Michaelis-Menton relation between [nicotine] and ΔF, with a Hill coefficient of 1. Other assumptions: $F_0$, 1000, counts; $EC_{50}$, 7, µM; $\Delta_{Fmax}$, 16000, counts. Concentration step, 0.1, µM

A, Unmodified dose-response relation on linear axes. Conditions as in the experiment of Figure 6A. The relation intersects the Y-axis at $F_0$.

B, as in A, overlaid with red axes denoting $F'_0 = 4F_0$, corresponding to 1.6 µM nicotine (or an unidentified endogenous ligand).

C, Simulation of the experiment of Figure 6A. The plot shows the dose-response relation near the origin of the red overlay. The X-axis, [nicotine]', represents incremental amounts of [nicotine]. The Y-axis, $\Delta F'/F'_0$ , represents incremental fluorescence.

| **Step** | Backbone restraint force-constant (kJ mol$^{-1}$ nm$^{-2}$) | Side-chain restraint force-constant (kJ mol$^{-1}$ nm$^{-}$) | Nicotine heavy atom restraints (kJ mol$^{-1}$ nm$^{-}$) | Time step (fs) | Simulation Time (ns) |
|---|---|---|---|---|---|
| 1 | 400 | 40 | 400 | 1 | 0.5 |
| 2 | 200 | 20 | 200 | 1 | 0.5 |
| 3 | 20 | 20 | 20 | 1 | 1 |
| 4 | 20 | 20 | 20 | 2 | 2 |

**Table S1**

Protocol for gradually releasing the restraints during MD simulations of the 10 nicotine-iDrugSnFR3a complexes, during the search for nicotine docking poses.

| Ranking | Mutation | Energy Score |
|---|---|---|
| 1 | R467W | -7.11 |
| 2 | D72Y | -6.77 |
| 3 | D72F | -6.28 |
| 4 | N39W | -5.97 |
| 5 | P475W | -5.95 |
| 6 | P79W | -5.79 |
| 7 | R36F | -5.51 |
| 8 | E24W | -5.28 |
| 9 | P328W | -5.28 |
| 10 | R36Y | -5.18 |
| 11 | P79H | -5.14 |
| 12 | N20W | -4.82 |
| 13 | P79M | -4.78 |
| 14 | N46W | -4.69 |
| 15 | Q431Y | -4.60 |
| 16 | K499W | -4.54 |
| 17 | P79F | -4.47 |
| 18 | N39F | -4.42 |
| 19 | R36W | -4.42 |
| 20 | K342W | -4.41 |
| 21 | D72H | -4.35 |
| 22 | E24Y | -4.32 |
| 23 | P400W | -4.19 |
| 24 | E24F | -4.13 |
| 25 | P328Y | -4.13 |
| 26 | N39Y | -4.13 |
| 27 | E513W | -4.11 |
| 28 | E24L | -4.11 |
| 29 | S325W | -4.03 |
| 30 | P79L | -3.99 |
| 31 | P475F | -3.98 |
| 32 | P507A | -3.96 |
| 33 | P475Y | -3.96 |
| 34 | D453W | -3.88 |
| 35 | N20Y | -3.85 |
| 36 | K454Y | -3.84 |
| 37 | R36I | -3.82 |

| | | |
|---|---|---|
| 38 | R52W | -3.75 |
| 39 | P400Y | -3.75 |
| 40 | E64L | -3.73 |
| 41 | P328F | -3.73 |
| 42 | R467Y | -3.73 |
| 43 | S325F | -3.72 |
| 44 | E476W | -3.65 |
| 45 | Q431F | -3.62 |
| 46 | N497Y | -3.61 |
| 47 | N497W | -3.58 |
| 48 | R467L | -3.55 |
| 49 | N497F | -3.52 |
| 50 | T413W | -3.52 |
| 51 | R36V | -3.48 |
| 52 | P79Y | -3.48 |
| 53 | E78W | -3.47 |
| 54 | S325Y | -3.44 |
| 55 | R36L | -3.43 |
| 56 | S9Y | -3.42 |
| 57 | K336W | -3.41 |
| 58 | P77W | -3.39 |
| 59 | K454W | -3.38 |
| 60 | P323W | -3.37 |
| 61 | T413Y | -3.35 |
| 62 | D439M | -3.34 |
| 63 | E27L | -3.34 |
| 64 | M418Y | -3.33 |
| 65 | P79I | -3.32 |
| 66 | E78Y | -3.31 |
| 67 | P323F | -3.31 |
| 68 | K454F | -3.27 |
| 69 | P475L | -3.27 |
| 70 | P464I | -3.26 |
| 71 | L384W | -3.24 |
| 72 | E24H | -3.24 |
| 73 | P400F | -3.23 |
| 74 | E517W | -3.21 |
| 75 | P475H | -3.21 |
| 76 | D72W | -3.20 |

| | | |
|---|---|---|
| 77 | Q431W | -3.19 |
| 78 | H68W | -3.19 |
| 79 | N46L | -3.19 |
| 80 | P79T | -3.16 |
| 81 | P350I | -3.15 |
| 82 | K342F | -3.15 |
| 83 | K51W | -3.14 |
| 84 | D434V | -3.13 |
| 85 | P464V | -3.13 |
| 86 | E78F | -3.12 |
| 87 | K362W | -3.11 |
| 88 | D501I | -3.11 |
| 89 | R341W | -3.07 |
| 90 | P79Q | -3.06 |
| 91 | D452H | -3.05 |
| 92 | E429W | -3.04 |
| 93 | P350W | -3.04 |
| 94 | R467F | -3.04 |
| 95 | P507T | -3.02 |
| 96 | N497V | -3.02 |
| 97 | P400L | -3.01 |
| 98 | N20F | -2.97 |
| 99 | D453H | -2.97 |
| 100 | K414W | -2.97 |

Table S2: Top 100 mutation outputs from single mutant stability analysis on iNicSnFR3a (Ranked by energy score)

| Ranking | Position | Mutation | Energy Gain |
|---|---|---|---|
| 1 | K342 | F | -4.51 |
| 2 | R52 | K | -3.98 |
| 3 | Q431 | Y | -3.96 |
| 4 | K51 | I | -3.85 |
| 5 | K454 | R | -3.84 |
| 6 | N20 | D | -3.56 |
| 7 | R467 | K | -3.56 |
| 8 | E476 | S | -2.88 |
| 9 | N46 | L | -2.74 |
| 10 | E27 | L | -2.04 |
| 11 | D452 | P | -2.04 |
| 12 | T413 | I | -1.95 |
| 13 | D453 | S | -1.85 |
| 14 | E64 | D | -1.73 |
| 15 | N497 | A | -1.73 |
| 16 | K414 | I | -1.61 |
| 17 | M418 | D | -1.59 |
| 18 | S325 | F | -1.43 |
| 19 | R36 | I | -1.42 |
| 20 | S9 | T | -1.11 |
| 21 | N39 | H | -0.67 |
| 22 | E517 | R | -0.58 |
| 23 | E78 | Y | -0.24 |
| 24 | E24 | - | 0.00 |
| 25 | K499 | - | 0.00 |
| 26 | E513 | - | 0.00 |
| 27 | K336 | - | 0.00 |
| 28 | D439 | - | 0.00 |
| 29 | L384 | - | 0.00 |
| 30 | D434 | - | 0.00 |
| 31 | K362 | - | 0.00 |
| 32 | D501 | - | 0.00 |
| 33 | R341 | - | 0.00 |
| 34 | E429 | - | 0.00 |

Table S2: Mutation outputs from sequence design on iNicSnFR3a on 34 previously identified 34 positions (Ranked by energy gain)
